# A functional network target for tic reduction during thalamic stimulation for Tourette Syndrome

**DOI:** 10.1101/2021.08.27.457924

**Authors:** Juan Carlos Baldermann, Christina Hennen, Thomas Schüller, Pablo Andrade, Veerle Visser-Vandewalle, Andreas Horn, Till A. Dembek, Jan Niklas Petry-Schmelzer, Joshua Niklas Strelow, Hannah Jergas, Jens Kuhn, Michael T. Barbe, Daniel Huys

## Abstract

**Background:** Deep brain stimulation (DBS) of the medial thalamus is an evolving therapy for severe, treatment-refractory Tourette syndrome (TS). It remains unanswered which functionally connected networks need to be modulated to obtain optimal treatment results.

**Methods:** We assessed treatment response of 15 patients with TS untergoing thalamic DBS six and twelve months postoperatively using the Yale Global Tic Severity Scale (YGTSS) tic score. For each time point, functional connectivity maps seeding from stimulation sites were calculated based on a normative functional connectome derived from 1000 healthy subjects. Resulting maps were analyzed in a voxel-wise mixed model for repeated measurements to identify patterns of connectivity associated with tic reduction.

**Results:** Connectivity of stimulation to the medial frontal cortex, bilateral insulae and sensorimotor cortex was associated with tic reduction. Connectivity with the temporal lobe, cerebellum, ventral striatum and orbitofrontal cortex was negatively associated. The overall connectivity pattern was robust to leave-one-out cross-validation, explaining 25 % of outcome variance (R = 0.500; p = 0.005).

**Conclusions:** We delineated a functional connectivity profile seeding from stimulation sites associated with TS-DBS outcome. This pattern comprised areas linked to the processing of premonitory urges and tic execution, thereby extending our current understanding of effective neuromodulation for TS.

## Introduction

Deep brain stimulation (DBS) for Tourette syndrome (TS) is an emerging therapy for severely affected, treatment-refractory patients. The scarce data from randomized controlled studies [1–3] along with results from an international registry [4] and a meta-analysis [5] suggest that thalamic DBS can be effective in reducing tics. However, firstly, there is still ambiguity about the ideal target for DBS in TS, and secondly, there are no reliable predictors of treatment response for this invasive procedure. Furthermore, little is known about the neural mechanisms that underlie the influence of DBS on tic symptomatology.

Previous studies have shown that structural connectivity profiles of DBS can predict clinical outcomes in patients with Parkinson’s disease [6], obsessive compulsive disorder [7] and more recently TS [8]. Such a structural connectomic approach allows the discrimination of fiber pathways that are associated with the respective DBS-induced clinical changes. Connectivity estimates derived from resting-state functional magnetic resonance imaging (fMRI) may extend these findings by deciphering polysynaptic, functionally connected networks that are not necessarily structurally linked. Thus, pursuing this connectomic DBS approach on a functional level, we performed an analysis of stimulation-dependent resting-state functional connectivity estimates to derive a network that, if modulated by thalamic DBS, explains tic improvement in DBS for severe TS.

## Methods

Fifteen patients (12 male) were retrospectively enrolled in the study (see supplemental data for demographics). This study is a secondary analysis of two clinical trials (drks.de, number DRKS00005316 and ClinicalTrials.gov, number NCT03958617) approved by the Ethics Committee of the University of Cologne. Written informed consent was provided by each patient. All subjects had been diagnosed with TS and underwent bilateral thalamic DBS surgery, receiving two quadripolar DBS electrodes (model 3389 [n = 11] or 3387 [n = 4]; Medtronic, Minneapolis, MN). In the majority of patients, the centromedian-parafascicular complex/nucleus ventro-oralis (Cm-Voi) was targeted (n = 12), while electrodes in the other patients were placed in the ventral vanterior and ventral lateral nucleus of the thalamus (VA/VL) with the most distal contacts residing in the field of Forel/subthalamic area (n = 3) (sFigure 1). Tic severity was assessed before DBS surgery as well as six and twelve months postoperatively with the tic score in the Yale Global Tic Severity Scale (YGTSS) (range 0-50). Over time, stimulation parameters had been adapted to reach the best clinical effect.

DBS electrodes were localized and volumes of tissue activated (VTAs) were calculated using Lead-DBS software (https://www.lead-dbs.org/) as described by Horn et al. [9]. In brief, postoperative CT scans were linearly coregistered to preoperative MRI images via Advanced Normalization Tools (ANTs; http://stnava.github.io/ANTs/). Subcortical refinement was applied to correct for brain shift that can occur due to brain surgery [9]. Images were normalized to the MNI 2009b nonlinear asymmetric template space using the SyN approach implemented in ANTs with additional subcortical refinement as implemented in Lead-DBS. DBS electrodes were localized by identifying the electrode artifacts in the postoperative image using the PaCER algorithm [10]. Afterwards, the electric fields were modeled with a finite element method based on an adaptation of the FieldTrip-SimBio pipeline integrated into Lead-DBS (https://www.mrt.uni-jena.de/simbio/; http://fieldtriptoolbox.org) based on the respective stimulation parameters as described elsewhere [6]. Binary VTAs were determined using a threshold of 0.2 V/mm.

Subsequently, functional connectivity between the bilateral VTAs and each brain voxel was calculated. To this end, we employed a publicly available normative resting-state-based functional connectome that was calculated from imaging data of 1,000 healthy participants from the Brain Genomics Superstruct Project (https://dataverse.harvard.edu/dataverse/GSP) [11], as described before [6]. For each pair of electric fields (from six- and twelve-months follow-up), the averaged functional connectivity across the 1,000 subjects to each voxel in a map of grey matter provided in SPM 12 was calculated by sampling time series from voxels within the electric fields in each subject of the normative connectome and correlating these time series with those of every other voxel. This resulted in a stimulation-dependent functional connectivity map that depicted the functional connectivity of each stimulation site in each patient. These maps (N = 30) were then analyzed employing a voxel-wise linear mixed model for repeated measurements, using the subject variable’s intercept as random factor and the functional connectivity estimate as fixed factor. The resulting connectivity map was thresholded at a significance level of p < 0.05. To validate the resulting model, we performed a leave-one-out cross-validation. To this end, we recalculated the described model, leaving one patient (one stimulation map for each of the two follow ups) out per time. Subsequently, we computed the similarity between the left-out patient’s stimulation-dependent connectivity maps (both time points) and the resulting contrast images of the YGTSS tic reduction variable from the full factorial design of the remaining sample (n = 28) by performing a spatial Spearman correlation between the individual stimulation map and the group model of optimal connectivity. Finally, we performed a linear regression of these similarity estimates with the empirical outcome, i.e. tic reduction after six or twelve months of chronic DBS. An overview of the methodological pipeline is given in figure 1.

**Figure 1:**
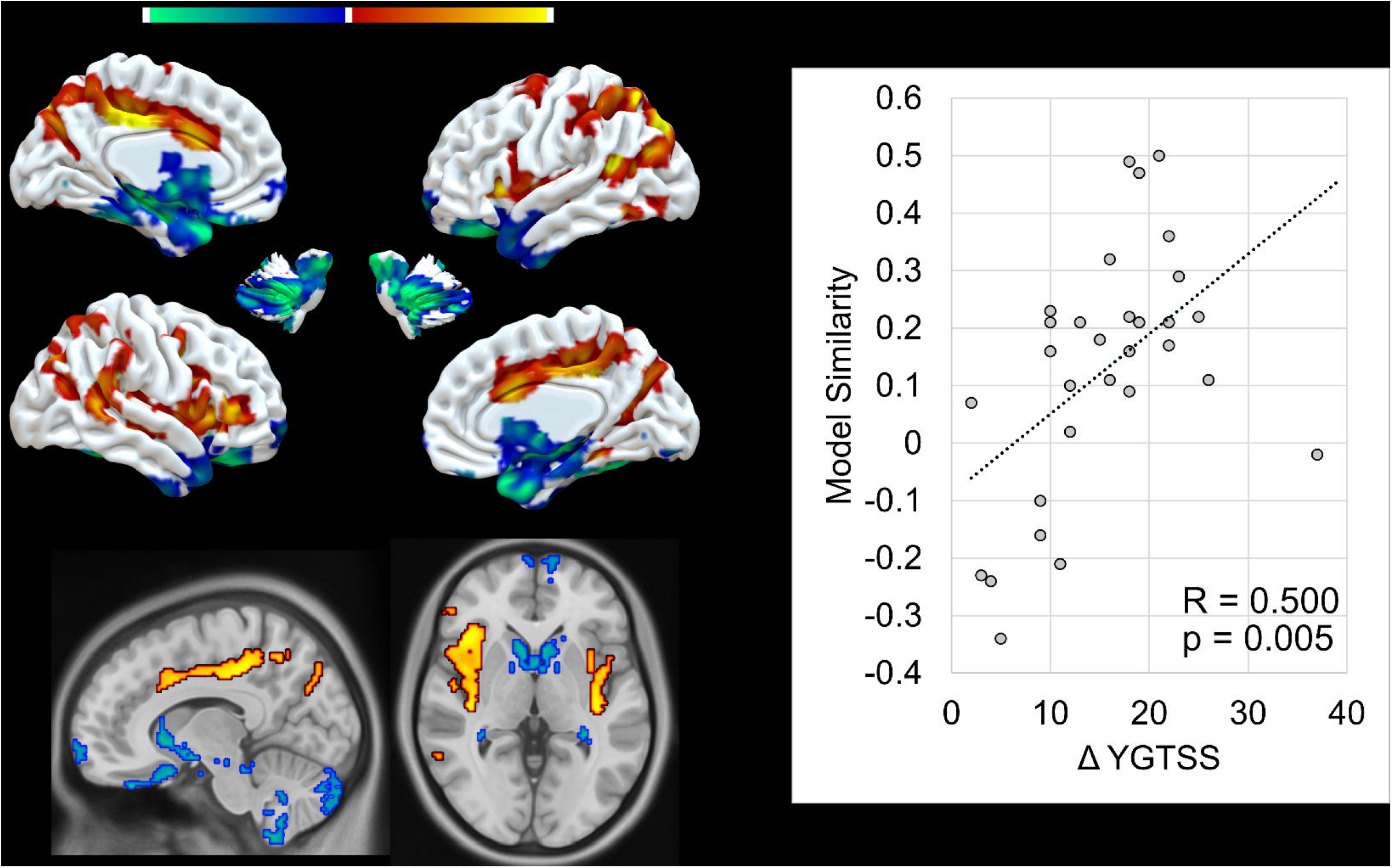
Overview of methodological workflow. A: For each bilateral VTA, the averaged functional connectivity to each grey matter voxel was calculated by sampling time series from voxels within the electric fields in each subject of a normative connectome derived from resting-state functional magnetic resonance imaging (rs-fMRI) and correlating these time series with those of every other voxel. B: This resulted in individual stimulation-dependent connectivity map, where each voxel contained a correlation coefficient, indicating positive or negative correlations with VTAs. C: Subsequently, we assigned the clinical outcome six and twelve months after DBS to each subject’s connectivity map. D We then calculated a voxel-wise mixed model for repeated measurements to determine the influence of functional connectivity estimates on the clinical outcome, resulting in a map of T-values, where higher values indicated a strong association of functional connectivity with tic reduction. E: Last, we validated the resulting model by performing a leave-one-out cross-validation.

## Results

Friedman’s test revealed a significant tic reduction from baseline (Median (M) = 42, interquartile range (IQR) = 18) to six (M = 23, IQR = 12) (z = 4.2; p < 0.001) and twelve months follow-ups (M = 23, IQR = 14) (z = 4.0; p < 0.001). Tic reduction was associated with functional connectivity to a cluster in the medial frontal cortex, covering the anterior and middle cingulate cortex and reaching the supplementary motor area. Likewise, connectivity of stimulation sites with the bilateral insula and primary motor and sensory cortices was associated with tic improvement. A negative association was observed for connectivity with the cerebellum, bilateral temporal cortex, orbitofrontal cortex and the ventral striatum. The overall pattern was robust to leave-one-out cross-validation, explaining 25 % of outcome variance (R = 0.500; p = 0.005) (Figure 2).

**Figure 2:**
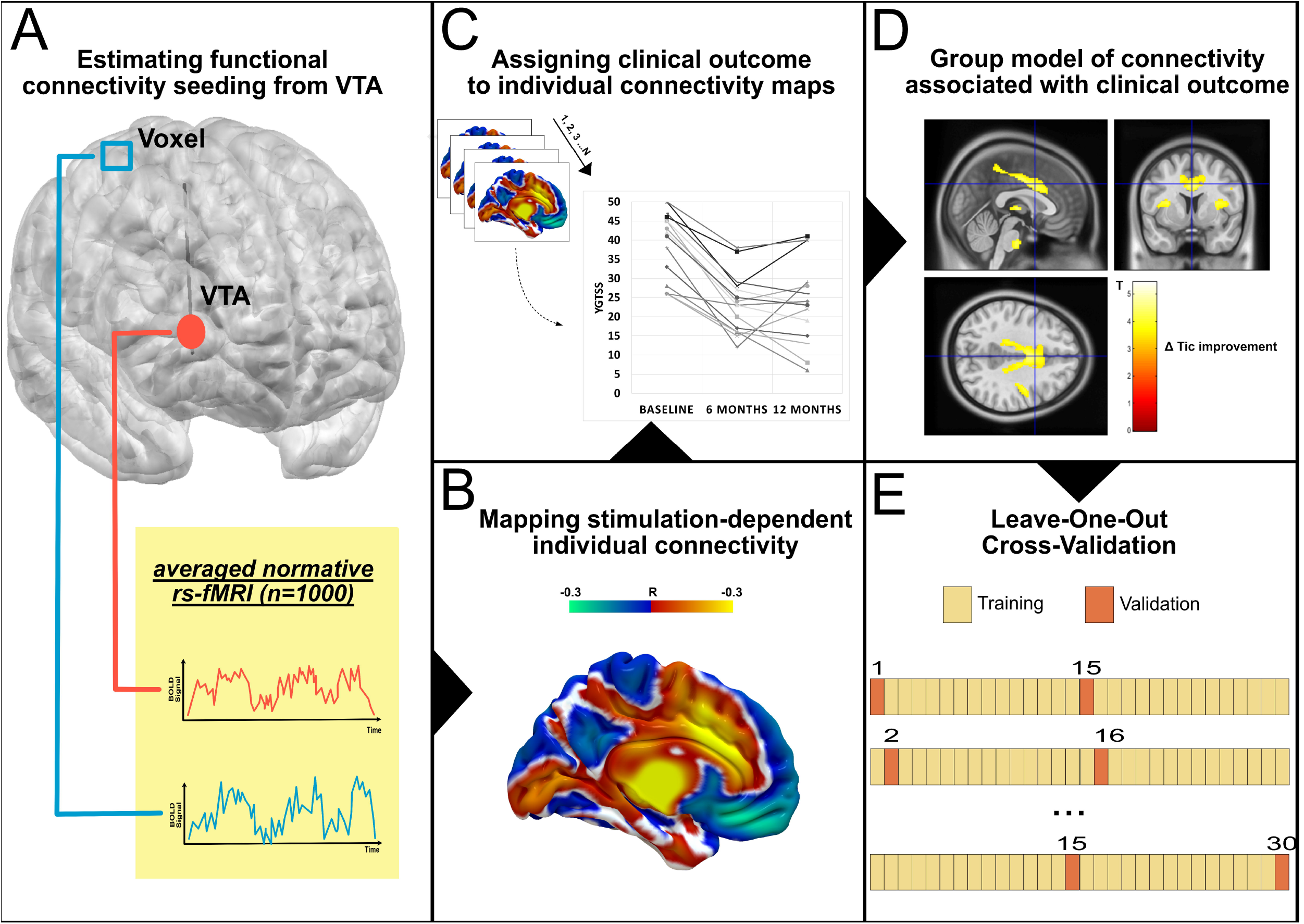
A: The final model of functional connectivity associated with tic reduction. Warm colors indicate higher T-values, thus a positive association of tic reduction and stimulation-dependent functional connectivity. Cool colors indicate a negative relationship. B: Illustration of the peak clusters. The strongest association was observed for a cluster in the medial frontal cortex, extending from the cingulate cortex to the ventral supplementary motor area and for bilateral clusters in the insulae. C: The final-model was validated, by applying a leave-one-out cross-validation. To this end, the model was recalculated repeatedly, each time leaving one subject out. The individual connectivity map of the left-out case was then compared with the model of the remaining subjects. The similarity index was then used to predict the clinical outcome (Change in the Yale Global Tic Severity Scale (YGTS)) in a linear regression, resulting in a significant relationship (R = 0.500; p = 0.005).

## Discussion

In this study, we delineated a functional connectivity profile of thalamic DBS for severe TS that is associated with tic improvement. We observed a pattern of clusters encompassing the medial frontal cortex and bilateral insulae, as well as sensorimotor cortices, related to tic reduction six to twelve months after DBS. This model of optimal stimulation-dependent connectivity was able to predict individual outcomes in a leave-one-out cross-validation.

Our findings add to very recent observations that stimulation-dependent structural connectivity profiles are associated with beneficial response of DBS for TS. Both applying normative or patient-specific [8] structural connectivity [12] revealed that fibers connecting to primary motor, premotor and primary sensory area are associated with of tic reduction after DBS. Our current approach of applying normative functional connectivity complements these studies, allowing to characterize polysynaptic networks relevant for DBS in TS.

The observed results are highly congruent to the studies based on structural connectivity [8,12] but reveal the importance of additional regions. The strongest association with tic reduction was found for connectivity with the bilateral insulae and the medial frontal cortex encompassing the cingulate cortex and supplementary motor area. Of note, the supplementary motor area, insula and cingulate area are active just before tic execution, followed by thalamic activation and lastly tic execution [13]. In most adult patients with TS, tics are preceded by a bothersome sensory phenomenon, the premonitory urge. This premonitory urge has been closely linked to activation in somatosensory cortices and the insula before tic onset [13,14]. Additionally, functional hyperconnectivity between insula and supplementary motor area is associated with increased perception of premonitory urge [15]. On a structural level, thinning of the insula was reported to correlate positively with severity of premonitory urges [16] while thinning of premotor areas correlated with tic severity [17,18]. Moreover, the supplementary motor area exhibits increased structural connectivity with the thalamus in patients with TS, which again correlates with tic severity [17].

Hence, the observed network associated with tic reduction can be linked to regions critically involved in premonitory urge and tic generation. It has been pointed out before that targeting the premonitory urge might be a promising approach for the treatment of TS by preventing tic learning and execution [8]. To date, it is unclear if thalamic DBS affects the premonitory urge to tic. Based on our results, we assume that thalamic DBS interferes with a sensorimotor network involved in both processing of the premonitory urge and tic genesis. However, further studies are warranted to decipher how exactly DBS may interrupt the cascade of aversive sensory urge phenomena and motor tic execution.

There are relevant limitations to our study: First, the sample size is restricted, due to the rarity of cases with severe TS treated with DBS. To partly overcome this obstacle, we included stimulation parameters from different time points to increase the power of our analysis, while statistically controlling for repeated measures. Furthermore, we used a normative connectome to calculate stimulation-dependent connectivity maps. The advantages and disadvantages of using patient-specific or normative connectivity data have been intensively discussed elsewhere, supporting the general notion that normative connectivity data is similarly capable of predicting treatment response in DBS [20]. For the case of severe and treatment-refractory TS, obtaining high quality MRI without sedation as necessary for resting-state fMRI is a major obstacle. Thus, we argue that relying on normative functional connectivity data can be a valuable option in these patients. Last, we applied a simplified model of the spread of the electric field and resulting VTAs. While this approach has been successfully used in a variety of similar publications [e.g. 5,20–23], more complex models [25,26] may yield even more accurate results in the future.

To conclude, we found that a specific functional connectivity profile seeding from the stimulation site was able to predict tic reductions after DBS. This pattern comprised areas strongly involved in the processing of the premonitory urge and genesis of tics, including the medial frontal cortex and bilateral insulae. Our results may help to increase our understanding of the neural mechanisms of neuromodulation in TS.

## Acknowledgment

JCB and VVV are funded by the Deutsche Forschungsgemeinschaft (DFG, German Research Foundation) – Project-ID 431549029 – SFB 1451. JNPS and TAD are supported by the Cologne Clinician Scientist Programe (CCSP), funded by DFG FI 773/15-1. A.H. was supported by the German Research Foundation (Deutsche Forschungsgemeinschaft, Emmy Noether Stipend 410169619 and 424778381 – TRR 295) as well as Deutsches Zentrum für Luft-und Raumfahrt (DynaSti grant within the EU Joint Programme Neurodegenerative Disease Research, JPND). A.H. is participant in the BIH-Charité Clinician Scientist Program funded by the Charité – Universitätsmedizin Berlin and the Berlin Institute of Health. JK received funding by the DFG (KFO-219, KU 2665/1-2) for the current study.

## Conflicts of Interest

MTB reports personal fees from Medtronic, Boston Scientific,Abbott (formerly St. Jude), GE Medical, UCB, Bial, IQWIG and grantsfrom Gondola, Felgenhauer-Stiftung and Forschungspool Klinische Studien, outside the submitted work.

