## Supplement for "A functional network target for tic reduction during thalamic stimulation for Tourette Syndrome"

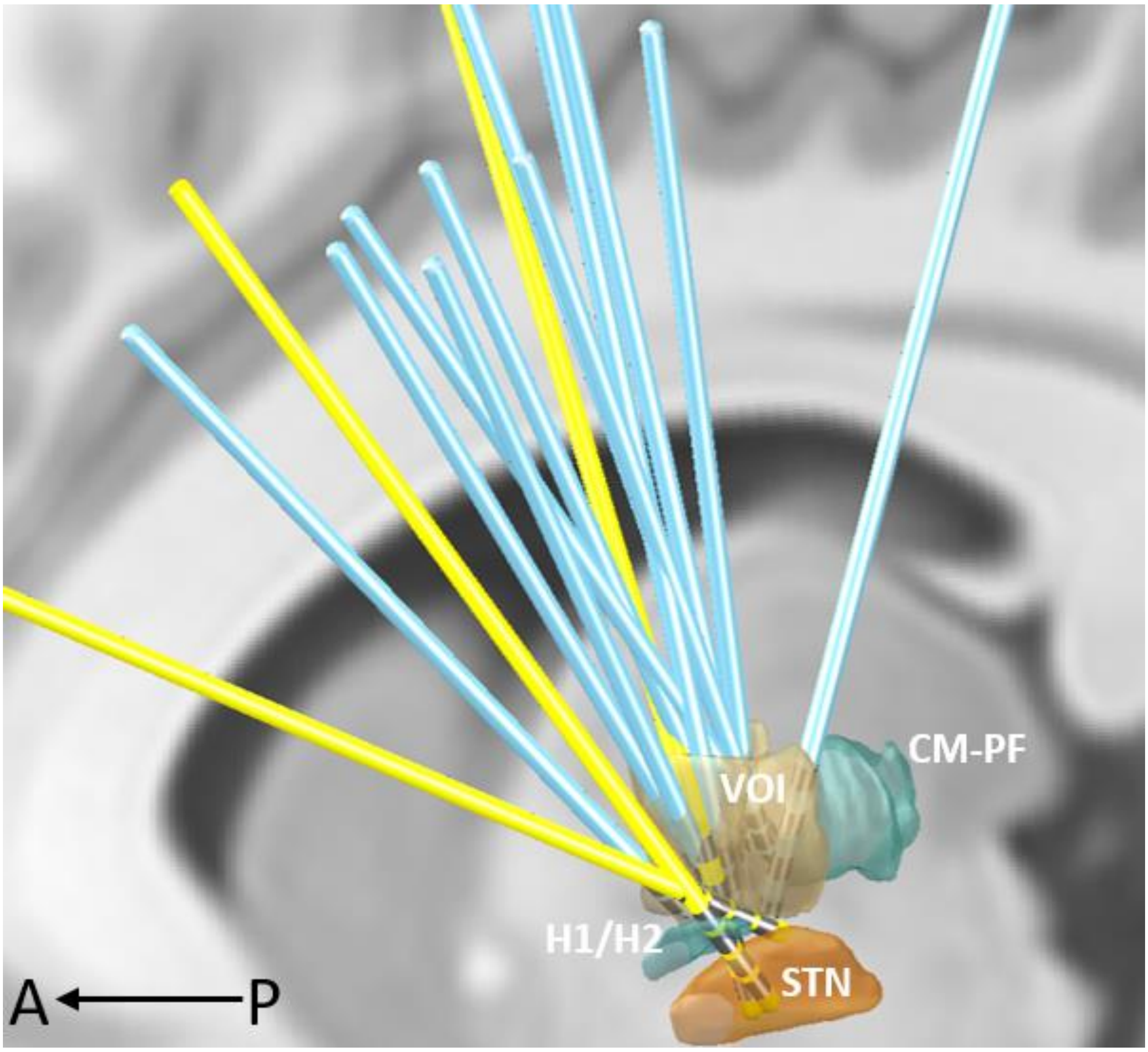


*SFigure 1: Visualization of electrode localization in standard MNI space. The target in three patients (yellow) was defined as the limbic part of the nucleus subthalamicus (STN) and field of Forel (H1 and H2) with the dorsal contacts residing in the nucleus ventro-oralis anterior and posterior (VOA/VOP). All other patients (blue) were targeted with the tip of the electrode in the centromedian-parafascicular complex and the nucleus ventro-oralis internus (VOI).*

*Table 1: Demographics*

| | **Mean/Ratio** | **Standard deviation** |
|---|---|---|
| **Females** | 3/15 | |
| **Age** | 33.2 | 12.4 |
| **Age at Tic onset** | 7.4 | 2.8 |
| **YGTSS Baseline** | 39.1 | 8.6 |
| **YGTSS Six months** | 23.3 | 7.4 |
| **YGTSS Twelve months** | 23.8 | 10.5 |

YGTSS = Yale Global Tic Severity Scale.
